# Enhancement of Therapeutic Transgene Insertion for Murine Phenylketonuria

**DOI:** 10.1101/2025.02.24.639678

**Authors:** Michael A Martinez, Daelyn Y Richards, Shelley R Winn, Adrian M Baris, Anne Vonada, Sandra Dudley, Laura Harper, Cary O Harding

## Abstract

Low *in vivo* transgene integration frequency limits the therapeutic efficacy of homology-directed repair (HDR)-mediated gene insertion as a treatment for Mendelian disorders. This study demonstrates improved efficacy of HDR-mediated gene insertion for the treatment of murine phenylalanine hydroxylase (PAH) deficiency, a model of human phenylketonuria (PKU), through pharmacologic inhibition of competing DNA repair pathways. Targeted integration of a *Pah-*expressing transgene into the hepatocytes of neonatal mice was enhanced with vanillin, a potent inhibitor of non-homologous end joining (NHEJ). This was further improved following combination of vanillin and novobiocin, an inhibitor of microhomology-mediated end joining (MMEJ). Combined NHEJ and MMEJ inhibition yielded PAH-expressing transgene insertions in approximately 10% of targeted alleles and was associated with a 70.6% decrease in serum phenylalanine. Demonstrating that pharmacologic inhibition of DNA repair pathways that compete with HDR can significantly enhance HDR-mediated transgene insertion *in vivo*.

## Introduction

Significant progress has been made in developing gene addition and gene correction methods for the treatment of inherited disorders. However, current gene addition techniques, which most frequently rely upon transient expression from viral episomes, offer only temporary relief due to their limited stability in dividing cells^1^. Meanwhile, gene correction methods that target single-nucleotide variants, such as base editors, remain impractical for treating genetic diseases associated with hundreds or thousands of pathogenic variants.

Phenylketonuria (PKU) is one such genetic disorder that exemplifies these challenges. PKU results from variations in the phenylalanine hydroxylase (*PAH*) gene that cause liver PAH enzyme deficiency, leading to neurotoxic accumulation of phenylalanine in the bloodstream. To date, over 1,500 unique variants have been associated with PAH deficiency in humans, with many individuals being compound heterozygous for two different *PAH* variants, challenging the clinical utility of gene correction tools that target a single variant. Therefore, targeted insertion of a complete functional *PAH-*expressing transgene into the liver genome is an attractive alternative, offering the potential to treat individuals harboring any *PAH* variants using a single therapeutic approach. To meet this need, researchers have made clever alterations to existing CRISPR/Cas9 machinery to create tools, such as PASTE editors, that are capable of large insertions^2,3^. However, while these inventions represent an exciting development for creating targeted insertions, they have not demonstrated robust *in vivo* editing success.

Alternatively, gene insertion through homology-directed repair (HDR) is a method that largely relies upon the endogenous repair machinery of the cell to incorporate a desired edit. A full expressing transgene is integrated into the genome by creating a targeted double-strand break (DSB) while simultaneously providing a repair template with the desired transgene flanked by regions of homology to the cut site. Repair of the DSB using the homologous recombination (HR) DNA repair pathway permanently incorporates the repair template into the genome. We previously demonstrated that pharmacological inhibition of a competing repair pathway, non-homologous end joining (NHEJ), significantly enhances the efficacy of HDR-mediated gene editing when correcting the *Pah^enu2^* missense variant^4^. In the present study, we demonstrate that this strategy can facilitate the integration of a large (>2,300 bp) murine *Pah* expression cassette resulting in a significant reduction of serum phenylalanine (Phe) concentration in a mouse model of PAH deficiency (Fig. 1). Subsequently, we observed that the simultaneous pharmacologic inhibition of two separate DSB repair pathways, NHEJ and microhomology-mediated end joining (MMEJ), resulted in a 70.6% reduction in serum Phe concentration when combined with our whole gene integration strategy. To our knowledge, this represents the most complete phenotypic correction of murine PAH deficiency achieved through HDR-mediated transgene insertion to date.

**Figure 1.**
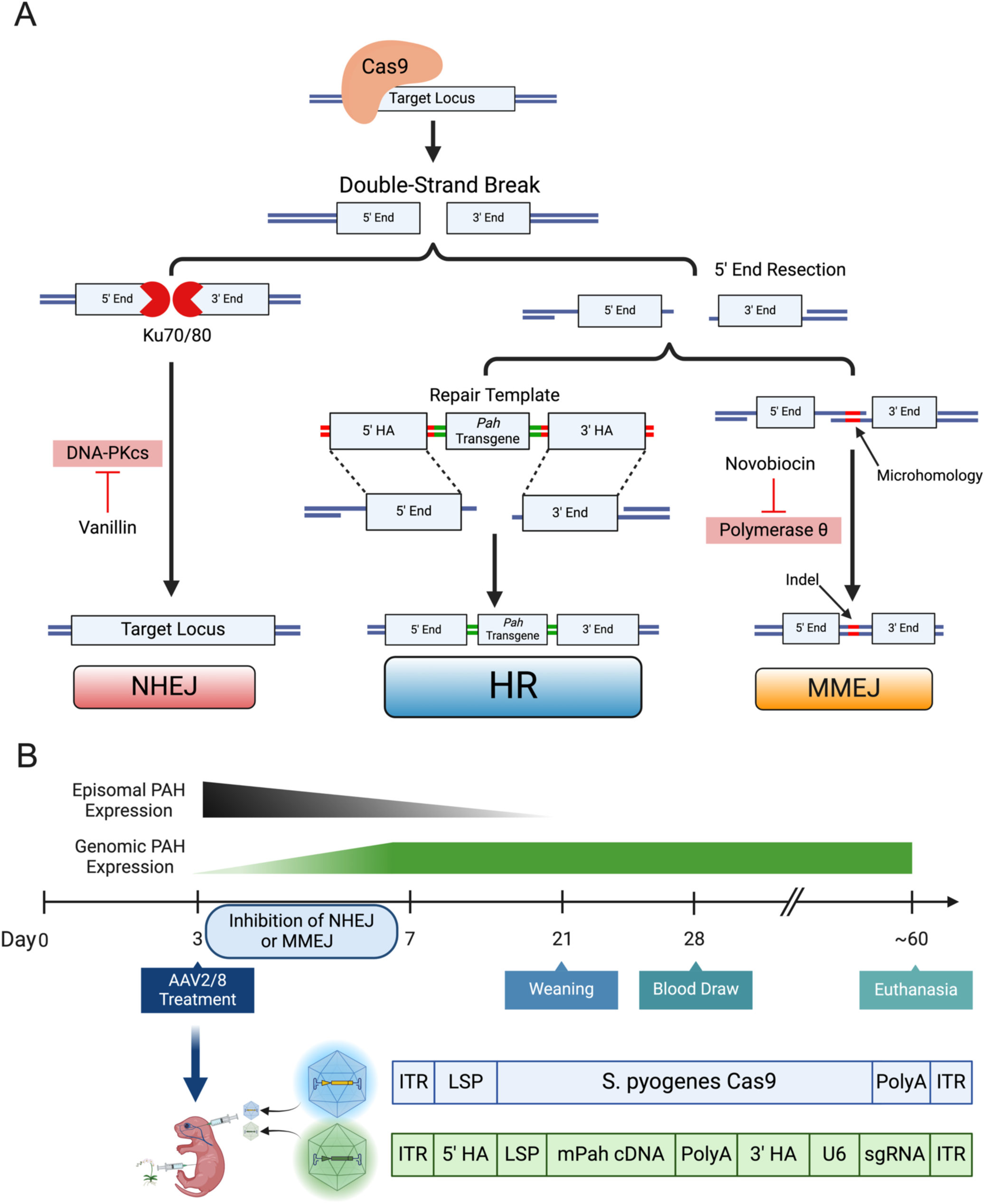
Experimental Design for HR-mediated Gene Editing Treatment of Murine PAH Deficiency. (**A**) Canonical model of DSB repair by non-homologous end joining (NHEJ), homologous recombination (HR), and microhomology-mediated end joining (MMEJ). Inhibition of DNA-PKcs and Polymerase theta, vital factors for NHEJ and MMEJ, is hypothesized to increase frequency of repair through HR. (**B**) Neonatal mice at neonatal day 3 (P3) receive dual AAV8 vectors through facial vein injection and daily IP injections of NHEJ or MMEJ inhibiting small molecules. Dual AAV8 treatment is composed of a streptococcus pyogenes Cas9 (spCas9) vector expressed from a liver specific promoter (LSP), and a repair template with 5’ and 3’ homology arms (HA) for integration into target genomic locus. Single-guide RNA (sgRNA) is delivered on the repair template, therefore requiring transduction with both vectors to induce a double-strand break (DSB). Episomal expression of murine PAH (mPAH) dissipates during hepatocyte turnover, while expression of repair template after integration into the genome remains.

## Results

### Pharmacologic inhibition of NHEJ promotes *Pah* transgene insertion into the PAH-deficient mouse liver genome

Previous work in our laboratory demonstrated significant reduction in serum Phe concentrations in the *Pah*^enu2/enu2^ mouse model. This was achieved through the co-administration of a recombinant adeno-associated virus (AAV) serotype 8 vector (AAV8) expressing *Streptococcus pyogenes* Cas9 (SpCas9) and a second AAV expressing a single guide RNA (sgRNA) and carrying a repair template designed specifically for the correction of the *Pah*^enu2^ missense variant in *Pah* exon 7^4^. However, this approach proved effective only in neonates that received injections of vanillin, an inhibitor of the NHEJ pathway^5^, to promote DNA repair through HDR (Fig. 1A)^4,6^. We hypothesized that vanillin treatment would similarly promote whole transgene insertion *in vivo*.

To test this hypothesis, three-day old *Pah^Δexon1/Δexon1^*mice (hereafter Dexon1 mice)^7^ received two separate AAVs (hereafter dual-AAV) via facial vein injection at a 1:1 ratio: one expressing SpCas9 and another carrying the repair template (4.2 × 10¹³ vg/kg total dose). Intraperitoneal (IP) injections of 100 mg/kg vanillin accompanied dual-AAV injection and were administered daily for five days following AAV delivery (Fig. 1B). The repair template vector delivered a 2,379 bp murine *Pah* expression cassette that was flanked by two 864 bp sequences with homology to *Pah* exon 1 (Fig. 2A).

**Figure 2.**
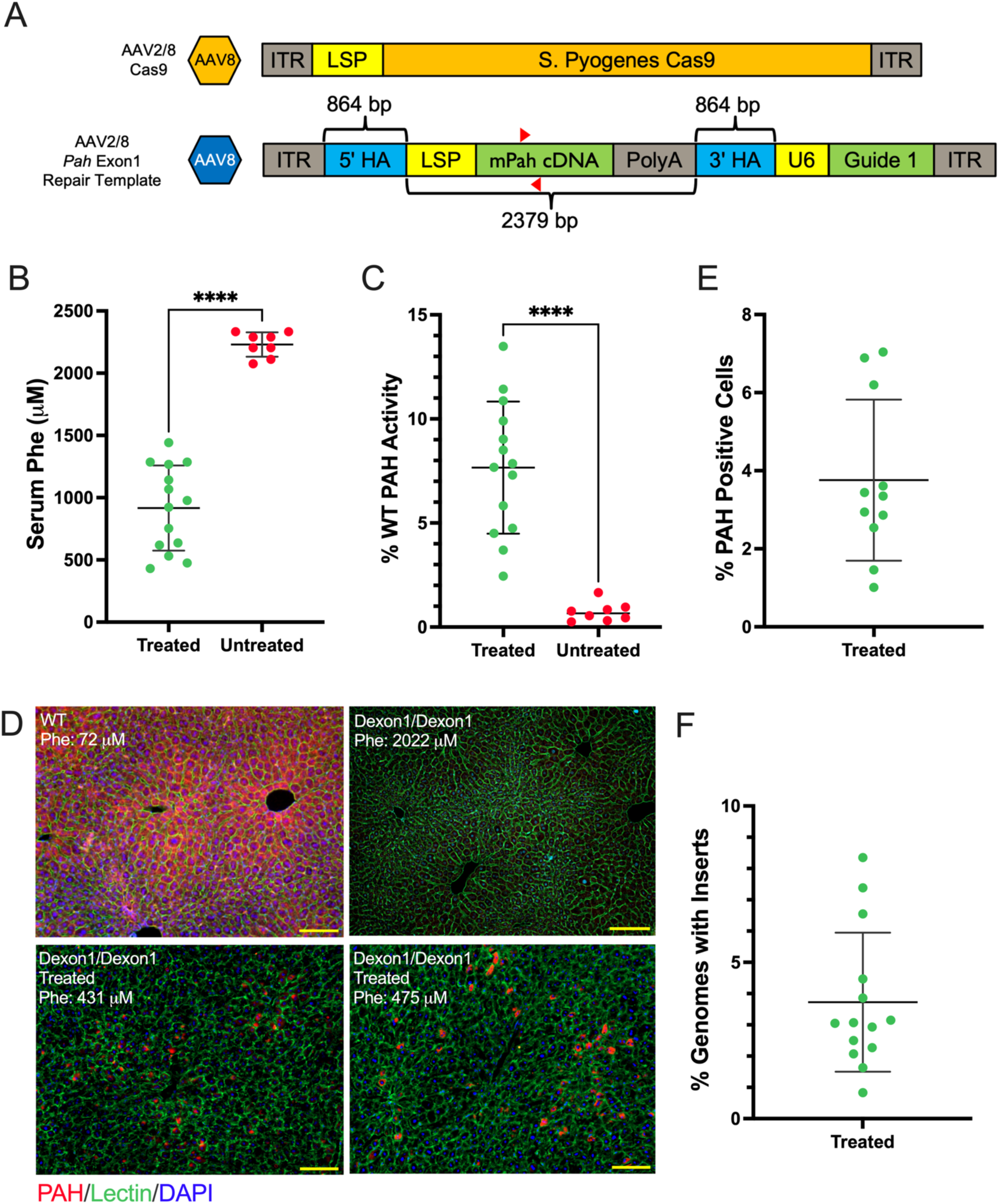
Vanillin Assisted Insertion of Pah Transgene in Murine Model of PKU. (**A**) Schematic of dual-AAV vectors including 5’ and 3’ homology arms (HA), liver specific promoter (LSP) including a bikunin enhancer and human thyroxine binding globulin (TBG) promoter, murine Pah cDNA, HGH poly(A) signal (PolyA), U6 promoter (U6), and sgRNA guide for targeted Cas9 cleavage (Guide 1). Red arrows indicate forward and reverse primers for qPCR. (**B**) Blood Phe concentrations in Pah*^Δexon1/Δexon1^*mice (n = 14) treated with dual AAV8 Cas9 and Pah Repair Template (mean = 917 ± 342 μM) versus control mice treated exclusively with AAV8 Pah Repair Template (mean = 2,122 ± 163 μM). All animals received facial vein injections of viral vectors (8.2 × 10^13^ vg/kg, 1:1 ratio) at day P3 with subsequent intraperitoneal injections of 100 mg/kg vanillin for 5 days following viral transduction. (**C**) PAH enzyme activity in treated (mean = 7.66 ± 3.17%) versus control (mean = 0.72 ± 0.45%) animals measured as percent of wild-type PAH activity. (**D**) Anti-PAH (Red, cytoplasmic), anti-lectin (Green, cell membrane), and DAPI (Blue, nuclear) of livers of treated (Bottom) and untreated (Top-left) Pah*^Δexon1/Δexon1^* animals as well as untreated C57BL/6 mice (Top-right). Scale bars, 100 μm. (**E**) Percent of liver genomes containing the mPah cDNA delivered in our repair template, as determined by qPCR, was 3.91 ± 2.5%. (**F**) Percent of hepatocytes with *Pah* transgene (mean = 3.76 ± 2.07%). \*\**P* < 0.01, \*\*\**P* < 0.001, and \*\*\*\**P* < 0.0001 by unpaired T-test. Data are means ± SD.

Serum Phe concentrations were reduced in the treated cohort to 916 ± 342 μM (mean ± SD) in comparison to 2,122 ± 163 μM in untreated animals (*P* <0.0001) (Fig. 2B). At euthanasia, mean liver PAH activity in treated mice was 7.67 ± 3.17% that of wild-type mice, significantly increased from 0.72 ± 0.45% measured in the untreated group (*P* <0.0001) (Fig. 2C).

Unlike other PAH-deficient models that produce a mutant PAH protein product, the PAH-null Dexon1 model allows visualization of successful repair events by PAH immunostaining (Fig. 2D). Quantification of PAH-positive hepatocytes revealed that 3.76 ± 2.07% of hepatocytes expressed PAH in the treated cohort (Fig. 2E). As murine hepatocytes are often polyploid, it can’t be assumed that each PAH-positive hepatocyte represents a single edited allele. Therefore, we employed qPCR on bulk liver genomic DNA using junction gap primers between *Pah* exons 7 and 8 to estimate the percentage of genomes with *Pah* cDNA insertions, yielding a mean insertion rate of 3.91% per haploid genome (Fig. 2F). However, this reaction detects the presence of the *Pah* cDNA anywhere in the liver genome and is therefore agnostic to the genomic integration site.

To confirm integration into the target site, select DNA sequences flanking the targeted genomic cut site and sequences from the transgene were enriched from bulk liver genomic DNA using custom probes and then sequenced on an Illumina next-generation sequencing (NGS) platform. This analysis confirmed integration of the *Pah* cDNA transgene into *Pah* exon 1 in treated mice (Fig. S1A). Interestingly, we also detected a significant amount of reads that included the U6 promoter and sgRNA from the AAV8 repair template, sequences that would not have integrated into the liver genome with the expected HDR-directed event. Two possible alternate molecular events would explain this result. First, integration of the entire AAV genome into the Cas9-induced DSB through NHEJ-mediated repair. Second, AAV concatemers, composed of serial repeats of a viral genome, may have been used as the substrate for the HDR-mediated repair, leading to integration of multiple copies of the whole viral construct. Previous studies have also observed frequent insertion of concatemers when delivering a repair template via AAV^8^. Additionally, we observed a small number (<120 reads) of partial Cas9 sequences within the integrated expression cassette, suggesting that a hybrid concatemer of the repair template and Cas9 episome was integrated into the target site in rare events (Fig. S1B).

### HDR-mediated transgene insertion into *Pah* Exon 7

One of the major advantages of whole transgene insertion is the ability to design a single sgRNA and repair template that can be efficiently advanced through clinical trials and produced at scale for use in most afflicted individuals. However, the homology arms of the repair template used in Figure 2 were tailored specifically to the unique deletion in the Dexon1 mouse model, which lacks most of the first exon of wild-type *Pah*. To enhance the rigor of this approach, and to create a universal repair template applicable to all known murine models of PKU, we designed a new repair template. This construct utilizes the *Pah* exon 7-targeting sgRNA, previously used to correct the *Pah^enu^*^2^ mutation^4^, and includes 800 base pair homology arms (Fig. 3A). Treatment of neonatal Dexon1 mice with the exon 7 repair template and Cas9 nuclease yielded a reduction in serum Phe concentrations, measured at 7 weeks of age, to 896 ± 325 µM (Fig. 3B). This is significantly reduced in comparison to mice that received the repair template without Cas9 (2026 ± 232 µM, *P* <0.0001). Liver PAH activity increased to 6.57 ± 2.11% in contrast to 0.67 ± 0.12% measured in RT-only treated animals (*P* <0.0001) (Fig. 3C). PAH-positive hepatocytes were observed in liver sections (Fig. 3D), with 7.44 ± 2.49% of bulk liver genomes possessing the *Pah* transgene (*P* <0.0001) (Fig. 3E).

**Figure 3.**
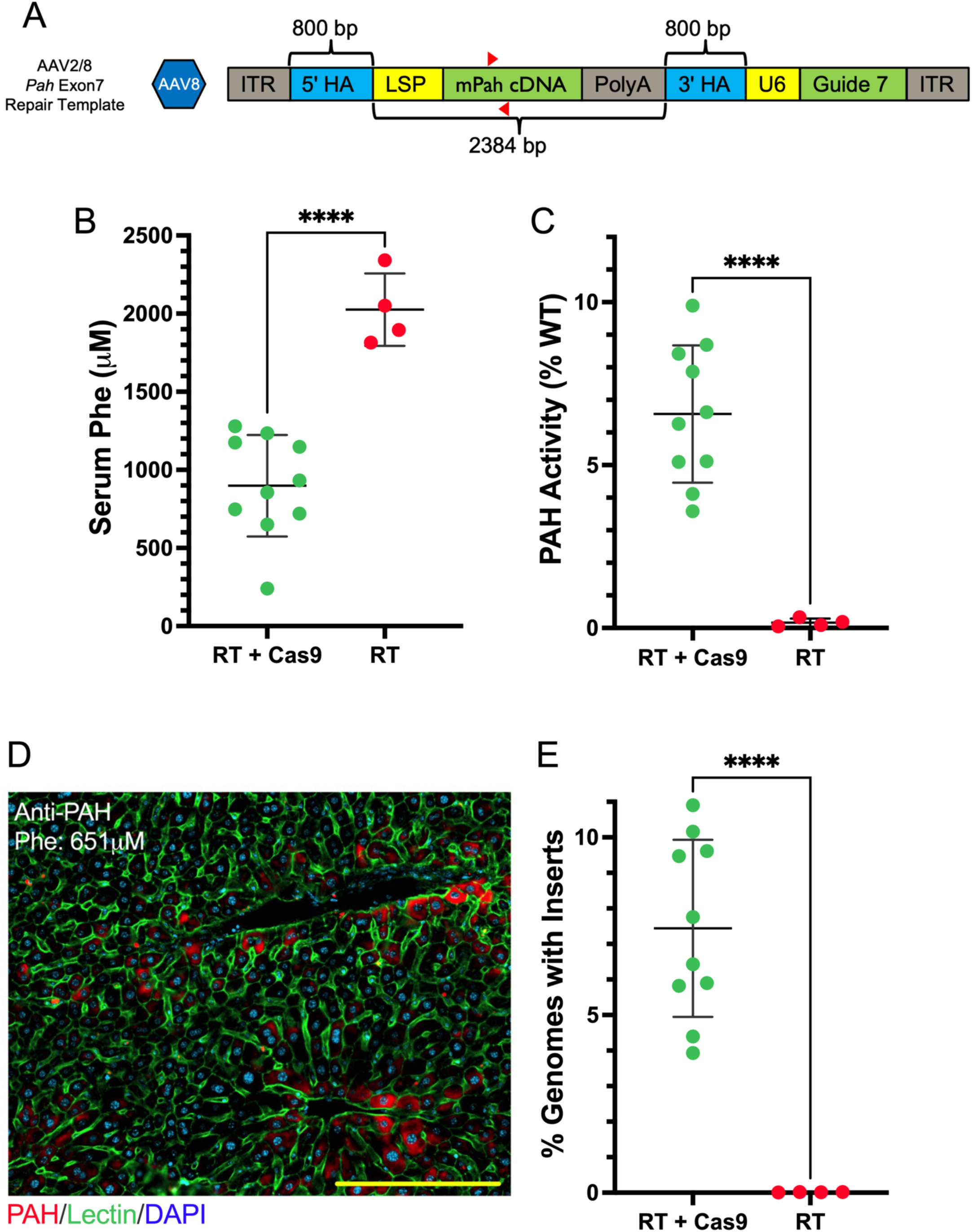
Insertion of the Transgene into *Pah* Exon 7. (**A**) Schematic of the repair template used to insert murine *Pah* cDNA into the *Pah* Exon 7 locus. Expression of sgRNA guide to this locus (Guide 7) under the expression of a U6 promoter (U6) and the adaptation of the 5’ and 3’ homology arms to the target locus are the major differences between Figure 1A. Animals were injected with a viral dose of 1.0 × 10^14^ vg/kg per vector and the same vanillin regime as in Figure 1. (**B**) Blood Phe concentrations in animals treated with both repair template (RT) and Cas9 carrying viral vectors (mean = 898 ± 324 μM, n = 10) versus those treated with only the RT (mean = 2026 ± 232 μM, n = 4). (**C**) PAH enzyme activity in RT + Cas9 treated animals (mean = 6.57 ± 2.11%) and RT treated animals (mean = 0.17 ± 0.12%). (**D**) Anti-Pah (Red, cytoplasmic), Anti-lectin (Green, cell membrane), and DAPI (Blue, nuclear) staining in liver section from RT + Cas9 treated animal. Scale bars, 100 μm. (**E**) Insertion frequency of the Pah cDNA (mean = 7.44 ± 2.49%) in treated animals, as determined by qPCR. \*\*\*\**P* < 0.0001 by unpaired T-test. Data are means ± SD.

AAV vector genomes form episomal concatemers that are capable of PAH expression without integrating into the genome^9,10^, but these episomes are unstable in dividing hepatocytes^11^. The persistent hyperphenylalaninemia in neonatally treated mice, given only the transgene-containing repair template, demonstrates three key points: (1) AAV episomes were largely lost during juvenile liver development, and physiologically-significant episomal PAH expression was not sustained into adulthood; (2) presence of homology arms in the repair template alone, without expression of a targeted nuclease, is insufficient for generating a clinically relevant number of genomic insertions, as has been previously shown^12,13^; and (3) although NHEJ-mediated insertions of fully intact AAV vector genomes into DSBs have been reported^14^, the frequency of these events was insufficient to correct serum Phe concentrations in Dexon1 mice treated with the repair template alone.

### Acetaminophen selection of Dexon1 hepatocytes harboring transgene insertions

To achieve a full phenotypic correction of Dexon1 mice, we sought to provide edited hepatocytes with a selective growth advantage. This would allow expansion of the edited hepatocyte population and increase their physiologic impact. Previous research has shown that the inclusion of a cytochrome P450 reductase (*Cypor*)-targeting short hairpin RNA (shRNA) can provide a selective advantage to edited hepatocytes in the presence of acetaminophen (APAP)-induced toxicity^15^. We hypothesized that addition of a *Cypor* shRNA to our transgene would promote the expansion of the PAH-expressing hepatocyte population during APAP-induced cell selection, resulting in further blood Phe reduction in PAH-deficient mice.

To test this hypothesis, we created a third repair template, still targeting *Pah* exon 7, with the addition of *Cypor* shRNA under transcriptional control of a ubiquitous U6 promoter (Fig. 4A). Neonatal Dexon1 mice were treated with this novel repair template, Cas9-expressing AAV, and vanillin as per prior experiments. At weaning, blood Phe concentrations remained extremely elevated (2128 ± 496 µM), suggesting low initial repair template integration frequency and liver PAH activity. Addition of 1.9% APAP (w/w) to the mouse chow was associated with a gradual decline in serum Phe (Fig. 4B), particularly in male mice. As a negative control, we treated another cohort of mice identically but did not provide APAP after weaning. Unlike our previous transgene insertion experiments, we observed a significant sex difference in this treatment response, with a mean genome insertion rate of 9.55 ± 1.16% in APAP treated male mice and 3.45 ± 2.48% in females (Fig. 4C). APAP-treated males achieved an average blood Phe concentration of 425 ± 205 μM (*P* <0.0001) while APAP-treated females achieved 1309 ± 603 μM (Fig. 4D).

**Figure 4.**
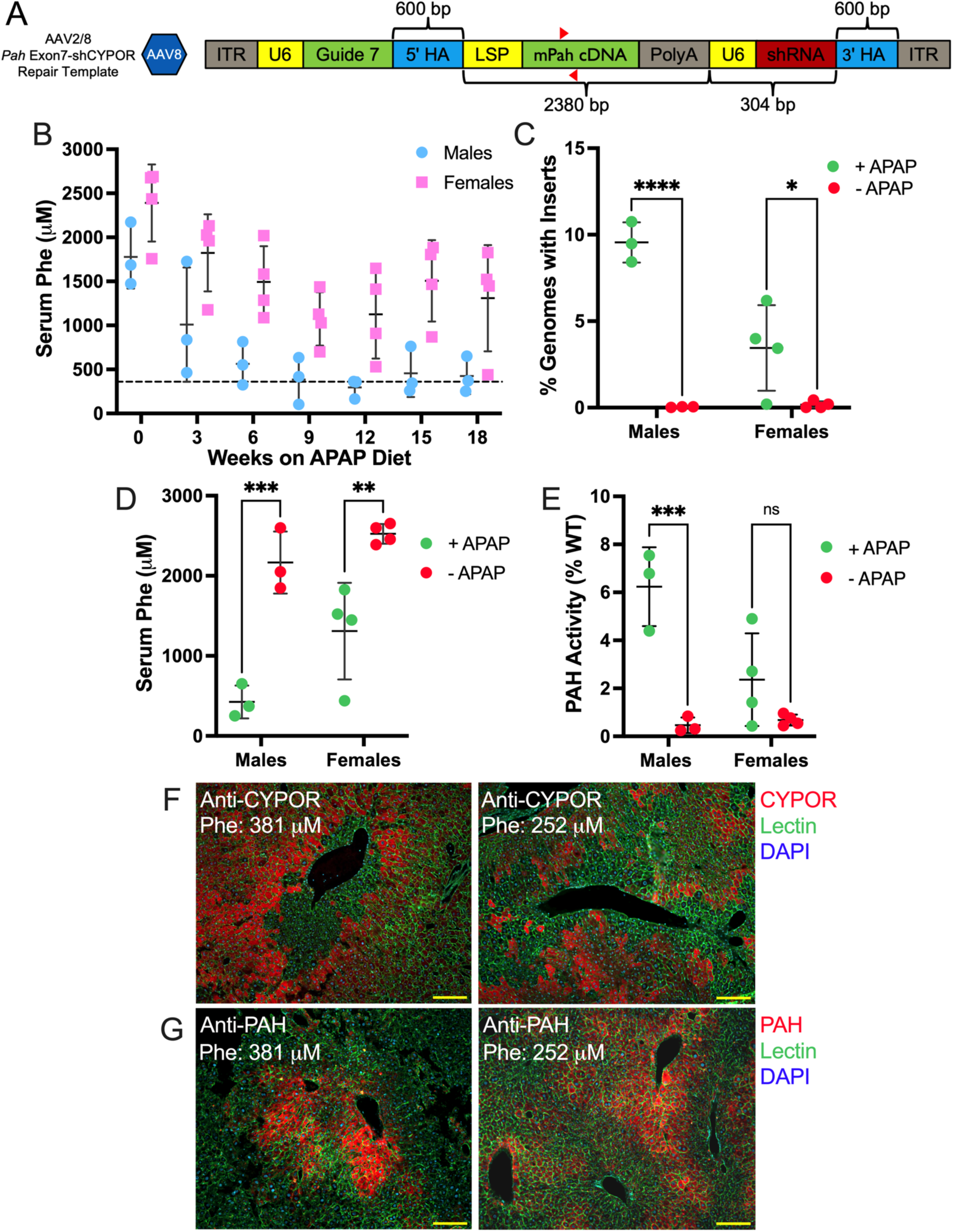
Providing *Cypor* shRNA Induced Selective Advantage to PAH Transgene under APAP Exposure. (**A**) Schematic of dual-AAV vectors including 5’ and 3’ homology arms (HA), liver specific promoter (LSP) including a bikunin enhancer and human TBG promoter, murine Pah cDNA, HGH poly(A) signal (PolyA), U6 promoter (U6), *Cypor*-targeting shRNA (shRNA), and sgRNA guide for targeted Cas9 cleavage (Guide 7). (**B**) Blood Phe concentrations over 18 weeks of APAP diet in male and female Dexon1 mice initiated after weaning. Dashed line indicates a therapeutic threshold of 360 μM. (**C**) Transgene insertion frequency in treated males that received APAP diet (9.55 ± 1.16%, n = 3) and those that did not (0.04 ± 0.02%, n = 3), and female that received APAP diet (3.45 ± 2.48%, n = 4) and those that did not (0.17 ± 0.20%, n = 4) (\**P* < 0.0193 and \*\*\*\**P* < 0.0001). (**D**) Terminal blood Phe concentrations between APAP treated males (425 ± 205 μM) and females (1309 ± 603 μM), as well as males (2165 ± 388 μM) and females (2524 ± 122 μM) that did not receive APAP selection but did receive viral vectors (\*\*\**P* <0.0005, and \*\**P* <0.0026). (**E**) PAH enzyme activity in treated males (6.24 ± 1.64%), treated females (2.36 ± 1.93%) and untreated males (0.46 ± 0.32%) and females (0.68 ± 0.23%) animals after 18 weeks of APAP induced hepatocyte selection (*P* < 0.1944 and \*\*\**P* <0.0006). (**F**) anti-CYPOR (Red, cytoplasmic), anti-lectin (Green, cell membrane), and DAPI (Blue, nuclear) in treated animals. (**G**) anti-PAH (red), anti-lectin (green), and nuclear stain (blue) in respective treated animals. Scale bars, 100 μm. Bonferroni’s multiple comparison test was used to calculate the significance of group differences. Significance calculated by two-way ANOVA with sex and treatment as variables. Data are means ± SD.

PAH enzyme activity in APAP-treated males measured 6.23 ± 1.64% of wild-type activity (*P* < 0.0001), while females exhibited mean activity of 2.36 ± 1.93% (Fig. 4E). Anti-CYPOR and anti-PAH immunohistology showed clear expansion of CYPOR deficient hepatocytes (Fig. 4F) as well as concomitant colonies of PAH positive cells in APAP-treated mice (Fig. 4G). Females with high blood Phe concentration demonstrated limited expansion of CYPOR deficient colonies (Fig. S2), suggesting that, while these mice did receive some low level of transgene integration, the founder population of edited cells failed to significantly expand under APAP selection.

### Inclusion of *Cypor* shRNA sequences alters AAV vector genome replication

Loss of AAV genome homogeneity and lower effective titers has been reported in vectors containing shRNAs^16^. DNA sequencing of the ‘*Pah* Exon7-shCypor’ (Fig, 4A) AAV vector stocks revealed that only 7.8% of *Pah* Exon7-shCypor vector genomes contained the desired construct while 61.8% of vectors consisted of so-called ‘snapback genomes’, which consisted of truncated AAV genomes without PAH experession elements^17^ (Figure. S3). The impaired production of fully functional AAV genomes in the presence of shRNA sequence is likely responsible for the low initial transgene integration frequency present prior to APAP selection.

### Severe motor disability observed in *Cypor* shRNA treated animals

Unexpectedly, animals treated with high dose AAV8 vectors carrying *Cypor* shRNA developed severe motor impairment, characterized by dystonic movements of all limbs, with a pronounced impact on the hindlimbs (Supplemental Movies 1 and 2). Out of 98 mice injected with *Cypor* shRNA-expressing AAV8 vectors, 53 exhibited motor dysfunction within 1-2 weeks post-treatment. Development of this adverse effect did not require co-administration of Cas9 vector. No animals receiving AAV8 repair templates lacking *Cypor* shRNA ever developed neurologic impairment, suggesting that *Cypor* shRNA expression is the underlying cause of this pathology. However, previous studies utilizing *Cypor* shRNA delivered to neonates via lentivirus vectors did not report such pathology^15^.

Administration of high AAV vector doses has been associated with abnormal histopathology of dorsal root ganglia but minimal clinical symptoms^18–20^. Also, injection of shRNA-expressing AAV vectors directly into brain has resulted in severe motor dysfunction, death, and neurodegeneration in several reported animal models^21–23^. However, gross anatomic examination of the brain, spinal cord, and DRG of our *Cypor* shRNA-treated mice revealed no clear pathology. Histologic examination (Nissl staining) of fixed brain sections from these mice likewise did not reveal any microscopic pathology (data not shown). Although AAV8 does not readily cross the blood-brain barrier (BBB) in adult mice, we considered the possibility that the BBB might be compromised in neonates. However, no AAV genomes were detected in brain tissue by qPCR in a limited sample of affected mice (n = 4). Therefore, the precise mechanistic cause of motor deficits in PAH-deficient mice treated with *Cypor* shRNA-expressing AAV8 remains undetermined.

### Simultaneous pharmacologic inhibition of NHEJ and microhomology-mediated end joining (MMEJ) synergistically enhances HDR-mediated gene insertion *in vivo*

Our previous research demonstrated that pharmacologic inhibition of NHEJ with vanillin produced a seven-fold increase of *in vivo* gene corrections as well as a corresponding increase in the number of small insertions and deletions (indels) at the targeted DSB^4^. We suspected that inhibition of NHEJ had increased the frequency of not only HDR, but also of microhomology-mediated end joining (MMEJ), which is inherently mutagenic^24,25^. High-throughput small molecule screens have identified the antibiotic Novobiocin (NVB) as a potent inhibitor of DNA polymerase theta, a key enzyme in MMEJ^26^. These screens also demonstrated that the coupling of NVB treatment with inhibitors of poly (ADP-ribose) polymerase 1 (PARP1), an essential NHEJ intermediate, leads to selective death of HR-deficient tumors^26^. This result indicates that inhibition of both MMEJ and NHEJ forces the cell to rely upon HR for DSB repair, a phenomenon that could be leveraged for increasing the efficacy of HDR (Fig. 1A). This hypothesis is supported by a recent report that used a potent inhibitor of DNA-PK catalytic activity (M3814) in the NHEJ pathway along with NVB to dramatically increase the efficiency of HDR for introduction of small mutations *in vitro*^27^.

To test this strategy, we evaluated simultaneous pharmacologic inhibition of NHEJ with vanillin, and of MMEJ with NVB, in a treatment strategy we have dubbed Vanbiocin. Five-day old Dexon 1 mice received vanillin, 100 mg/kg and NVB, 50 mg/kg by IP injection followed by facial vein injections of the AAV8 Cas9 and AAV8 Pah Exon 7 repair template. Vanbiocin treatment was repeated daily for two additional days (three days total). An initial treatment attempt in 3-day old mice with five-day total Vanbiocin treatment duration had been associated with lethargy, poor feeding, and death of all pups by day 10. Delaying treatment onset and shortening the duration of Vanbiocin therapy improved tolerability and survival (8 out of 11 treated pups). At six weeks of age, serum Phe concentrations in mice that had received both AAV8 vectors and Vanbiocin treatment measured 631 ± 211 μM (n = 8) in comparison to 2,134 ± 197 μM in untreated mice (Fig. 5A). Mean insertion frequency of the repair template at euthanasia in Vanbiocin treated mice was 9.71 ± 3.62% with mean liver PAH activity of 6.14 ± 1.15% of wild-type activity (Fig. 5B-C). Anti-PAH staining of liver sections showed colonies of PAH positive hepatocytes primarily near the central hepatic veins (Fig. 5D). Coat pigmentation, which is light brown in C57BL/6 models of PKU due to phenylalanine-mediated inhibition of melanin production, was rescued in mice treated with Vanbiocin (Fig. 5E).

**Figure 5.**
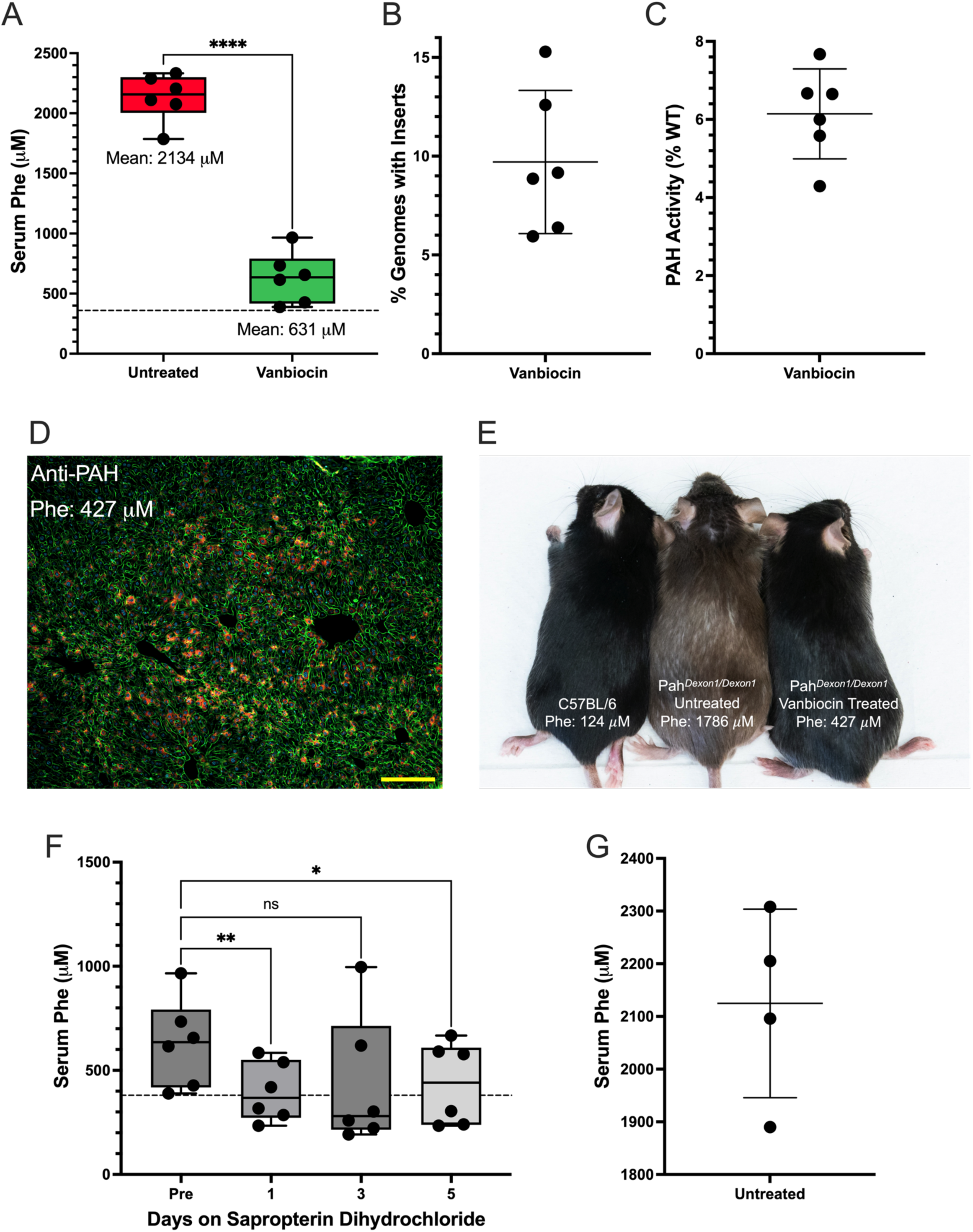
Inhibition of NHEJ and MMEJ Increases Efficacy of Homology-Directed Recombination *in vivo*. (**A**) Blood Phe concentrations in untreated mice (mean = 2134 ± 197 μM, n = 6) and animals treated with Cas9 and RT AAV vectors with Vanbiocin (Vanillin and Novobiocin) (mean = 631 ± 211 μM, n = 6). Black dashed line indicates a therapeutic threshold of 360 μM. Data are means ± SD. \*\**P* <0.0027 and \*\*\*\**P* <0.0001 by paired t-test. (**B**) Insertion frequency of the mPah cDNA (mean = 9.71 ± 3.62%) in treated animals, as determined by qPCR. (**C**) PAH enzyme activity in Vanbiocin treated mice (mean = 6.14 ± 1.15%) measured as percent of wild-type PAH activity. (**D**) anti-PAH (Red), anti-lectin (green), and nuclear stain (blue) in treated animals. Scale bars, 100 μm. (**E**) Coat color rescue of wild-type C57BL/6, untreated Dexon1 mice, and Vanbiocin treated mice. Baseline blood Phe concentrations without sapropterin provided. (**F**) Blood Phe concentrations in Vanbiocin-treated animals was measured over a 5-day course of daily treatment with 100 mg/kg sapropterin dihydrochloride, administered via oral gavage 6 hours prior to retro-orbital blood collection. Black dashed line indicates a therapeutic threshold of 360 μM. *P* <0.1188, \**P* <0.0242, and \*\**P* <0.0037 by one-way ANOVA. (**G**) Blood Phe concentrations in the progeny (mean = 2124, n = 4) of Vanbiocin treated female post-weaning. Dunnett’s multiple comparison used with all ANOVAs. Data are means ± SD.

Female PAH deficient mice, if untreated and hyperphenylalaninemic, are unable to produce viable offspring due a severe maternal PKU effect^28^. Here, gene insertion/Vanbiocin-treated Dexon1 dams successfully gestated and reared pups through weaning while consuming normal chow and without additional Phe-lowering treatment. All progeny from matings between Vanbiocin-treated Dexon1 dams and untreated homozygous Dexon1 sires exhibited severe hyperphenylalaninemia at weaning, indicating that all offspring lacked liver PAH activity. All pups from this pairing were homozygous for the Dexon1 mutation, indicating that genomic insertions in the dams were liver specific and had not been transmitted to germ cells (Fig. 5G).

### Hyperphenylalaninemia in Vanbiocin-treated Dexon1 mice is responsive to sapropterin supplementation

PAH deficiency in humans is represented by a broad spectrum of disease severity that is determined in an individual by the specific *PAH* variants inherited and the resultant amount of residual liver PAH activity. Individuals with 2-5% residual liver PAH activity may exhibit mild hyperphenylalaninemia (blood Phe ≤ 600 µM) while consuming an unrestricted diet. In many of these individuals, treatment with pharmacologic doses of tetrahydrobiopterin^29^, the natural pterin cofactor for PAH^30^, or its pharmaceutical analog, sapropterin dihydrochloride^31^, augments PAH activity and lowers blood Phe. We hypothesized that since Vanbiocin-treated Dexon1 mice exhibited partial correction of serum Phe concentrations, sapropterin treatment may improve Phe metabolism in the mice and lead to further reduction in serum Phe concentrations. We provided gene-edited, Vanbiocin-treated animals with 100 mg/kg sapropterin via oral gavage, daily, over the course of 5 days, and observed a static reduction in serum Phe at 396 ± 142 μM (*P* = 0.0089), near the therapeutic threshold for PKU (Fig. 5F).

### Non-invasive assessment of *in vivo* Phe oxidation capacity in Vanbiocin-treated gene-edited Dexon1 mice using stable isotope loading

Following oral loading with 1-^13^C-L-phenylalanine and timed collection of breath CO_2_, we non-invasively documented restored total body Phe metabolism in Vanbiocin-treated gene-edited Dexon1 mice through measurement of ^13^CO_2_ excreted in breath (Fig. S5A). Plotting total ^13^CO_2_ production vs. final serum Phe concentration in Vanbiocin treated Dexon1 mice revealed an exponential curve (R^2^=0.9655) (Fig. S5B), in a relationship consistent with the known Michaelis-Menten enzyme kinetics for PAH with allosteric regulation by Phe^32^. No significant difference in Phe oxidation was detected in Vanbiocin treated animals with or without sapropterin supplementation. These results suggest potential clinical utility for stable isotope breath testing as a method for non-invasively assessing total body Phe oxidation in PKU clinical gene therapy trials, without the need for more invasive liver biopsy.

## Discussion

Our results demonstrate that, through small molecule inhibition of NHEJ and MMEJ, large transgenes can be inserted into the genome of animals at a physiologically relevant frequency using HDR-dependent editing. The results presented here demonstrate an increased *in vivo* editing efficacy over the gene correction strategy we previously described^4^, but with the advantage of inserting a full *Pah* transgene. If further refined, this treatment strategy could provide the same therapeutic relief as has been observed in AAV-mediated gene addition trials for humans with PKU^33–37^ with the advantage of being a long-term treatment that is not dependent upon of the survival of viral episomes. As we have presented, this technique is not overly dependent upon homology arm length or a unique locus, suggesting this form of HDR has the potential to treat, not only PKU, but also a wide variety of other liver-based genetic diseases.

To enhance the efficacy of HDR-mediated transgene insertion, we provided edited hepatocytes with a selective advantage conferred via shRNA inhibition of CYPOR. However, we have found that the presence of shRNA in a repair template dramatically inhibits HDR-mediated gene editing when delivered by AAV8 vectors by both dramatically reducing the effective viral titer and providing competitive viral isoforms in the form of ‘snapback’ genomes. Future research that could deliver shRNAs in viral vectors without inducing titer heterogeneity may be able to rescue the integration frequency we observed in our earlier experiments (Fig. 1,2), while conferring edited cells with a selective advantage that will require far less time and APAP toxicity to provide a therapeutic effect. Furthermore, expression of *Cypor* shRNA from AAV8 in neonatal mice appears toxic to the nervous system.

We found greater success through pharmacologic inhibition of NHEJ and MMEJ, the two DNA repair pathways that predominantly compete with the planned HDR event. Vanbiocin treatment resulted in mean transgene insertion frequency of almost 10% of hepatocyte genomes and 70.6% reduction in blood Phe concentration, which is to our knowledge the best efficacy from whole transgene insertion reported to date in PAH-deficient mice. Furthermore, sapropterin supplementation reduced mean blood Phe even further to 396 µM, just above the target lifelong treatment threshold for humans with PKU^38^. In humans with sapropterin-responsive PAH deficiency, sapropterin treatment may act through three possible molecular mechanisms: 1) stabilization of misfolded variant PAH monomers, 2) allosteric activation of PAH activity, or 3) rescue of *PAH* variants that alter the K_M_ for binding of the native tetrahydrobiopterin cofactor^39^. Given that Dexon1 mice do not produce any native PAH protein, the sapropterin supplement would only be acting on wild-type PAH expressed from the integrated transgenes: folding and kinetics of the transgene-expressed PAH protein was expected to be normal. Therefore, sapropterin responsiveness in Vanbiocin mice was unexpected. Sapropterin-mediated allosteric activation of transgene-expressed PAH is the likely mechanism to explain this observation. However, we were unable to demonstrate a significant increase in whole body 1-^13^C-Phe oxidation following sapropterin treatment. So, the mechanism of Phe lowering following sapropterin treatment in Vanbiocin mice remains unexplained. However, the results suggest that PAH-deficient humans exhibiting a partial response to clinical gene therapy may benefit from sapropterin treatment and experience further blood Phe reduction.

CRISPR/Cas9 nuclease treatment has been shown to produce off-target mutations across the genome in major regulatory segments and cause major translocation events due to aberrant repair^40^. However, one of the benefits of performing HDR in neonates, as we have done in this study, is that viral episomes responsible for Cas9 expression are rapidly lost during liver development, preventing long-term exposure to Cas9 and reducing the risk of genotoxicity^41–43^. In our previous work using the same *Pah* exon 7 sgRNA, we sought insertions and deletions at several of the most likely predicted off-target sites in genomic liver DNA and found these to be below the limits of detection for the amplicon based assay employed^4^. We have not repeated a search for off-target edits here, but a more exhaustive analysis for off-target genotoxicity will be necessary before this approach, with human genome directed reagents, could be advanced to the clinic.

HDR is dependent upon active cell division, as HR is active predominantly in late S/G2 phase of the cell cycle. Therefore, this treatment approach is maximally effective in neonatal or infantile animals with rapidly growing livers. Pharmacologic inhibition of NHEJ and MMEJ may not overcome the lack of HR activity in post-mitotic cells of adult animals; further studies are needed to understand whether pharmacologic enhancement of HR-mediated editing will be effective in adult animals.

While these promising results represent significant progress in HDR as a treatment option, in our opinion, we have not demonstrated sufficient efficacy to justify bringing this treatment design to human clinical trials. Individuals who are heterozygous for a *Pah* variant frequently exhibit a dominant-negative phenotype^30^, wherein mutant PAH monomers form tetramers with wild-type monomers, leading to reduced enzymatic activity. As such, a greater transgene insertion frequency may be necessary both to disrupt mutant PAH monomer production and to restore wild-type PAH activity, thereby lowering blood Phe into the normal physiologic range. Therefore, further optimization is necessary to consistently achieve robust disease correction in murine and larger animal disease models. Targeted inhibition of other proteins involved in non-HR repair pathways (PARP1/2 or 53BP1 for instance), promotion of factors required in HR (BRCA1/2, MRN, ATM), or artificially promoting cell-cycle turnover could enhance the frequency of site-specific gene insertion and improve the physiologic relevance of this treatment strategy. In conclusion, while significant progress has been made, realizing the full potential of HR-mediated whole gene insertion for therapeutic applications will require a deeper understanding of the molecular mechanisms involved as well as strategic interventions to enhance precision and efficiency.

## Materials and Methods

### Animal husbandry

Animal care and experimentation were performed in accordance with the guidelines of the Department of Comparative Medicine, Oregon Health & Science University, and the NIH Guide for the Care and Use of Laboratory Animals. C57BL/6-*Pah^Δexon1/Δexon1^* mice (hereafter Dexon1 mice), which are homozygous for a deletion of exon 1 in the murine *Pah* gene, lack production of any PAH monomer and are completely deficient in liver PAH activity^7^. *Dexon1* mice exhibit severe hyperphenylalaninemia on an unrestricted diet. Neonatal Dexon1 mice for these experiments were generated through breeding of *Pah^Δexon1/Δexon1^*sires to *Pah^Δexon1/+^* dams. All mice were fed tap water and standard mouse chow (LabDiet Picolab Rodent Diet 5LOD, St. Louis, MO, USA) *ad libitum*, with the exception of mice on APAP diet, providing approximately 24% protein and 1.04% L-Phe by weight, except the breeders that received high-energy chow (LabDiet Rodent High Energy Diet 5058, St. Louis, MO, USA), providing approximately 22% protein and 0.99% L-Phe. Given that adult mice consume approximately 5 g chow per day, daily L-Phe intake was estimated to be approximately 50 mg/day. The animals were housed under a standard 12-h:12-h on-off light cycle. All surgical procedures were carried out with inhaled isoflurane general anesthesia to minimize pain. Mice that exceeded 20% weight loss were given 2 days on a high fat diet to regain weight. Animals were euthanized through CO_2_ or isoflurane inhalation followed by cardiac puncture.

### Genotyping of *Dexon1* mice

To assess whether mice were homozygous for the Dexon1 deletion a PCR of Pah exon 1 was performed (5’-*GCAGGGAACAGTCATCCTTAAT*-3’ forward primer, 5’-*ACCAATGAGCCACCCAAAG*-3’ reverse primer)^44^. An allele with the Dexon1 deletion will yield a product of 498 bp, while a wild-type allele will yield a product of 757 bp.

### Stable isotope breath test

1-^13^C-L-Phenylalanine (Cambridge Isotope Laboratories, Tewksbury, MA) was diluted to 5 mg/ml in sterile H_2_0. Each mouse received 100 μl injections IP and placed in custom breath collection chamber with CO_2_ analyzer. Exetainer collection tubes (Labco, Lampeter, Ceredigion UK) were used to collect 5 ml of breath sample every five minutes from baseline to 30 minutes. CO_2_ was measured in ppm simultaneously with each breath collection. The concentration of radiolabeled CO_2_ was analyzed (Metabolic Solutions LLC, Nashua, NH) and used to calculate metabolic turnover of L-Phenylalanine as previously reported^45^. Data are reported as % 1-^13^C-L-phenylalanine dose metabolized.

### Vector design and viral production

Plasmids were designed in Snapgene (Version 7.0.1). Whole plasmid sequencing was performed by Plasmidsaurus using Oxford Nanopore Technology to confirm intact ITRs and correct plasmid structure. The AAV8 vectors used to express PAH and/or *Cypor* shRNA after insertion into *Pah* exon 7 in our study pAAV[Exp]-U6>sgRNAex7-5’HA and pAAV[Exp]-U6>gRNA-mPAH-Ex7 were constructed and packaged by VectorBuilder (Chicago, IL). Titers were estimated using qPCR-based quantification of ITR sequences. The vector IDs for these constructs were VB221216-1108hwe and VB220601-1003sae respectively, which can be used to retrieve detailed information about the vector on www.vectorbuilder.com.

The OHSU Molecular Virology Support Core produced large-scale preparations of recombinant AAV8 virus for the insertion of *Pah* into Exon 1, referenced as AAV8 *Pah* Exon1 Repair Template, using standard triple plasmid transfection procedures into cultured HEK293 cells and purified by iodixanol gradient ultracentrifugation.

### Delivery of Small Molecules and Viral Vectors

Vanillin was dissolved in saline (0.9% w/v) and heated in 50 ^°^C degree water bath with frequent vortexing. Novobiocin sodium salt (Thermo Fisher Scientific, Waltham, MA) was dissolved in 400 μL of DMSO and heated in a 50 ^°^C water bath before slowly adding 600 μL of saline preheated to 50 ^°^C. Viral vectors, diluted in saline, were injected into the facial vein of neonatal mice at either P3 or P5. An IP injection of the desired small molecule cocktail was performed immediately after facial vein injection and would continue for 3-5 days.

### Serum phenylalanine

Serum phenylalanine was determined using an established fluorometric protocol^46^.

### PAH enzyme activity

PAH enzyme activity was determined using a previously reported radioactive chromatography assay, with modifications^47^.

### Immunohistochemistry

Liver tissue was immediately placed in 4% paraformaldehyde after organ harvest and kept overnight. Tissue was then placed in a gradient of 10, 20, and 30% sucrose. Tissue was then frozen imbedded in OCT and 8–10-micron sections were collected using cyrostat (Tanner Scientific, Sarasota, FL) and transferred to slides (Superfrost Plus, Fisher, Pittsburgh, PA). Slides were permeabilized in 0.1-0.25% triton X-100, diluted in PBS, for 12 minutes, followed by a five-minute wash with PBS that was repeated three times. Blocking buffer of 10% normal goat serum (Sigma-Aldrich, Burlington, MA) was added to slide for 30 minutes. PAH antibody (Bioworld, BS3704) was diluted 1:500 in blocking buffer and incubated on slides overnight at 4 °C. Slides were again washed three times with PBS for five minutes. Alexa 594/555 goat-anti-rabbit antibody (Invitrogen, Eugene, OR) was diluted 1:500 in blocking buffer in addition to lectin conjugated antibody (Vector Laboratories, FL-1171) at the same dilution. Secondary antibodies were incubated for 2 hours at room temperature and washed three times with PBS for five minutes. If staining for CYPOR was desired anti-CYPOR antibody (Abcam: ab252084) was diluted 1:200 in blocking buffer and incubated overnight at 4 °C. Slides were washed 3×5 in PBS for a final time before Dapi-Fluoromount G (Southern Biotech) was added to sections and cover slips were mounted to the slides and sealed with nail polish. Slides were imaged using a Ziess ApoTome 2 microscope.

### Image analysis

Image analysis for the quantification of PAH positive hepatocytes (Fig. 2D) was performed in ImageJ.

### APAP preparation and delivery

To produce 1.9% (w/w) APAP chow, acetaminophen (6.84 g) was dissolved in 200 proof ethanol (50 ml). The APAP/ethanol solution was added to high fat chow (LabDiet Picolab Rodent Diet 5LOD, St. Louis, MO, USA) (360 g) and shaken until fully absorbed; the chow was then allowed to dry overnight in a fume hood. To induce selection of CYPOR-negative hepatocytes, gene edited mice were placed on 1.9% APAP treated chow for at least three weeks beginning after weaning. As APAP toxicity is associated with liver dysfunction which can artifactually elevate blood Phe, measurement of blood Phe concentrations was carried out in experimental animals only after APAP chow had been discontinued and standard chow substituted for at least one week prior to phlebotomy for measurement of blood Phe.

### Insertion Frequency via qPCR copy number assay

The percentage of haploid liver genomes containing PAH cDNA insertions in edited mouse livers was measured by real-time quantitative polymerase chain reaction (qPCR) using a primer set complementary to *Pah* exons 7 and 8, that spans an intron boundary and therefore will not amplify the genomic *Pah* gene in a PCR protocol with a brief amplification cycle. The primers used span exon-exon boundaries between exon 7 and 8, with the as 5’-*GCTGGCTTACTGTCGTCTC*G-3’ the forward primer and 5’-*CATGTCCCAAGAGTTCATGACAG*-3’ as the reverse primer, with a final product size of 136 bp. A standard curve was created by diluting a plasmid carrying *Pah* cDNA to calculate the insertion frequency of the transgene in treated liver samples.

### Target enrichment and NGS sequencing

Custom probes specific to the *Pah* expression cassette and 400 bp flanking the genomic cut site were produced (101001, Twist Biosciences, San Francisco, CA). Using the ‘Twist Target Enrichment Standard Hybridization v1’ protocol target sciences were enriched and read sequenced on Illumina NovaSeq 6000.

### NGS data analysis

NGS data was aligned to the murine genome, and other genomes of interest (see Supplementary Figure 1) using the QuasR pipeline (Version 1.44.0). Aligned data was visualized using the Integrated Genomics Viewer desktop application.

### AAV genome sequencing

Single stranded AAV sequencing was performed by Plasmidsaurus using Oxford Nanopore Technology with custom analysis and annotation shown in Supplemental Figure 1.

### Qualitative assessment of urinary phenylpyruvate excretion

During the stable isotope breath testing, urine samples were collected passively over 30 minutes by placing Whatman 3MM Chr filter paper on the floor of the breath collection chamber. Areas of fecal contamination on the filter paper were excluded by its yellow appearance on long wave UV illumination. Urine spots that appeared purple on UV illumination due to the presence of urea were excised from the filter paper and eluted with 2 ml 0.01 M NH_4_OH. Semi-quantitative organic acid analysis was performed by trimethylsilyl (TMS) derivitization followed by gas chromatography-mass spectrometry (GC-MS). In this assay, the concentration of the TMS derivative of phenylpyruvate was measured relative to the added internal standard, undecanedioic acid^48^. In this comparison, relative phenylpyruvate concentrations were not corrected for urine creatinine excretion.

### Graphical illustrations

Graphical illustration of DNA repair and experimental workflow shown in Figure 1 was created in www.biorender.com.

### Statistical analysis

Significance of Figure 2, Figure 3, Figure 4C, and Supplemental Figure A were analyzed by unpaired T-test. An Ordinary one-way ANOVA was used to analyze significant in Figure 4D, 4E, Figure 5A and 5B. The significance of supplemental Figure 3B was performed by one-way ANOVA. A post-hoc Dunnetts multiple comparison test to establish significant differences between groups. All statistical analysis was performed using Graphpad Prism version 10.2.2 for Mac (GraphPad Software, La, Jolla, California, www.graphpad.com). Full data and analysis results can be found in Supplmental Table 1.

#### Contributions

M.A.M. designed, executed, and analyzed data from experiments in this work, and wrote this manuscript with assistance from all authors. D.Y.R. designed and executed initial experiments targeting *Pah* Exon 1. S.R.W. performed facial vein injections, tissue processing, and performed PAH enzyme activity assessment in later experiments. A.M.B. performed NGS data analysis and visualization. A.V advised the experimental design and analysis for experiments utilizing Cypor shRNA-mediated selective advantage. S.D. and L.H. bred and genotyped Dexon1 mice. C.O.H. assisted in the design of all experiments in this manuscript as well as its writing.

## Abbreviations

HDR: homology-directed repair
PAH: phenylalanine hydroxylase
Phe: phenylalanine
PKU: phenylketonuria
NHEJ: non-homologous end joining
MMEJ: microhomology-mediated end joining
PASTE: programmable addition via site-specific targeting elements
DSB: double strand break
SpCas9: Streptococcus pyogenes Cas9 nuclease
sgRNA: single guide RNA

## Data availability statement

Datasets generated or analyzed during the current study are available from the corresponding author upon request.

## Acknowledgements

We thank Biomarin and Dr. Brian Whitlock at The University of Tennessee for providing the Kuvan (sapropterin dihydrochloride) used in this study. We thank the OHSU viral production core for AAV production and tittering. We also thank Aaron Bieleck for assistance photographing mouse coat color. Finally, we would like to thank Dr. Charles Venditti for providing equipment to execute ^13^CO_2_ breath testing.

## References

1. Daya, S. & Berns, K. I. Gene therapy using adeno-associated virus vectors. Clin. Microbiol. Rev. 21, 583–593 (2008).

2. Yarnall, M. et al. Drag-and-drop genome insertion of large sequences without double-strand DNA cleavage using CRISPR-directed integrases. Nat. Biotechnol. 41, (2023).

3. Böck D et al. In vivo prime editing of a metabolic liver disease in mice. Sci. Transl. Med. 14, (2022).

4. Richards, D. Y. et al. AAV-Mediated CRISPR/Cas9 Gene Editing in Murine Phenylketonuria. Mol. Ther. Methods Clin. Dev. 17, 234–245 (2020).

5. Durant, S. & Karran, P. Vanillins—a novel family of DNA-PK inhibitors. Nucleic Acids Res. 31, 5501–5512 (2003).

6. Paulk, N. K., Loza, L. M., Finegold, M. J. & Grompe, M. AAV-mediated gene targeting is significantly enhanced by transient inhibition of nonhomologous end joining or the proteasome in vivo. Hum. Gene Ther. 23, 658–665 (2012).

7. Richards, D. Y. et al. A novel Pah-exon1 deleted murine model of phenylalanine hydroxylase (PAH) deficiency. Mol. Genet. Metab. 131, 306–315 (2020).

8. Suchy, F. P. et al. Genome engineering with Cas9 and AAV repair templates generates frequent concatemeric insertions of viral vectors. Nat. Biotechnol. 43, 204–213 (2025).

9. Fisher, K. J. et al. Recombinant adeno-associated virus for muscle directed gene therapy. Nat. Med. 3, 306–312 (1997).

10. Grieger, J. C., Choi, V. W. & Samulski, R. J. Production and characterization of adeno-associated viral vectors. Nat. Protoc. 1, 1412–1428 (2006).

11. Nakai, H. et al. Extrachromosomal recombinant adeno-associated virus vector genomes are primarily responsible for stable liver transduction in vivo. J. Virol. 75, 6969–6976 (2001).

12. Porteus, M. H., Cathomen, T., Weitzman, M. D. & Baltimore, D. Efficient Gene Targeting Mediated by Adeno-Associated Virus and DNA Double-Strand Breaks. Mol. Cell. Biol. 23, 3558– 3565 (2003).

13. Jasin, M. Genetic manipulation of genomes with rare-cutting endonucleases. Trends Genet. TIG 12, 224–228 (1996).

14. Hanlon, K. S. et al. High levels of AAV vector integration into CRISPR-induced DNA breaks. Nat. Commun. 10, 4439 (2019).

15. Vonada, A. et al. Therapeutic liver repopulation by transient acetaminophen selection of gene-modified hepatocytes. Sci. Transl. Med. 13, (2021).

16. Xie, J. et al. Short DNA Hairpins Compromise Recombinant Adeno-Associated Virus Genome Homogeneity. Mol. Ther. 25, 1363–1374 (2017).

17. Zhang J et al. Subgenomic particles in rAAV vectors result from DNA lesion/break and non-homologous end joining of vector genomes. Mol. Ther. Nucleic Acids 29, (2022).

18. Hordeaux J et al. Adeno-Associated Virus-Induced Dorsal Root Ganglion Pathology. Hum. Gene Ther. 31, (2020).

19. Bevan, A. et al. Systemic gene delivery in large species for targeting spinal cord, brain, and peripheral tissues for pediatric disorders. Mol. Ther. J. Am. Soc. Gene Ther. 19, (2011).

20. Gombash, S. E. et al. Systemic Gene Delivery Transduces the Enteric Nervous System of Guinea Pigs and Cynomolgus Macaques. Gene Ther. 24, 640–648 (2017).

21. Martin, J. N. et al. Lethal toxicity caused by expression of shRNA in the mouse striatum: implications for therapeutic design. Gene Ther. 18, 666 (2011).

22. McBride, J. L. et al. Artificial miRNAs mitigate shRNA-mediated toxicity in the brain: Implications for the therapeutic development of RNAi. Proc. Natl. Acad. Sci. 105, 5868–5873 (2008).

23. Ehlert, E. M., Eggers, R., Niclou, S. P. & Verhaagen, J. Cellular toxicity following application of adeno-associated viral vector-mediated RNA interference in the nervous system. BMC Neurosci. 11, 20 (2010).

24. Seol, J.-H., Shim, E. Y. & Lee, S. E. Microhomology-mediated end joining: Good, bad and ugly. Mutat. Res. 809, 81–87 (2018).

25. Sinha, S. et al. Microhomology-mediated end joining induces hypermutagenesis at breakpoint junctions. PLoS Genet. 13, e1006714 (2017).

26. Zhou, J. et al. A first-in-class Polymerase Theta Inhibitor selectively targets Homologous-Recombination-Deficient Tumors. *Nat*. Cancer 2, 598–610 (2021).

27. Riesenberg, S. et al. Efficient high-precision homology-directed repair-dependent genome editing by HDRobust. Nat. Methods 20, 1388–1399 (2023).

28. Levy, H. L. & Waisbren, S. E. Effects of untreated maternal phenylketonuria and hyperphenylalaninemia on the fetus. N. Engl. J. Med. 309, 1269–1274 (1983).

29. Kure, S. et al. Tetrahydrobiopterin-responsive phenylalanine hydroxylase deficiency. J. Pediatr. 135, 375–378 (1999).

30. Kaufman, S. Tetrahydrobiopterin: The Metabolic and Molecular Bases of Inherited Disease. (The John Hopkings University Press, Baltimore, Maryland, 1997).

31. Levy, H. L. et al. Efficacy of sapropterin dihydrochloride (tetrahydrobiopterin, 6R-BH4) for reduction of phenylalanine concentration in patients with phenylketonuria: a phase III randomised placebo-controlled study. Lancet Lond. Engl. 370, 504–510 (2007).

32. Fitzpatrick, P. F. Allosteric Regulation of Phenylalanine Hydroxylase. Arch. Biochem. Biophys. 519, 194–201 (2012).

33. Oh, H.-J., Park, E.-S., Kang, S., Jo, I. & Jung, S.-C. Long-term enzymatic and phenotypic correction in the phenylketonuria mouse model by adeno-associated virus vector-mediated gene transfer. Pediatr. Res. 56, 278–284 (2004).

34. Mochizuki, S. et al. Long-term correction of hyperphenylalaninemia by AAV-mediated gene transfer leads to behavioral recovery in phenylketonuria mice. Gene Ther. 11, 1081–1086 (2004).

35. Ding, Z., Georgiev, P. & Thöny, B. Administration-route and gender-independent long-term therapeutic correction of phenylketonuria (PKU) in a mouse model by recombinant adeno-associated virus 8 pseudotyped vector-mediated gene transfer. Gene Ther. 13, 587–593 (2006).

36. Harding, C. O. et al. Complete correction of hyperphenylalaninemia following liver-directed, recombinant AAV2/8 vector-mediated gene therapy in murine phenylketonuria. Gene Ther. 13, 457–462 (2006).

37. Ahmed, S. S. et al. Sustained Correction of a Murine Model of Phenylketonuria following a Single Intravenous Administration of AAVHSC15-PAH. Mol. Ther. Methods Clin. Dev. 17, 568–580 (2020).

38. Smith, W. E. et al. Phenylalanine hydroxylase deficiency diagnosis and management: A 2023 evidence-based clinical guideline of the American College of Medical Genetics and Genomics (ACMG). Genet. Med. Off. J. Am. Coll. Med. Genet. 27, 101289 (2025).

39. Blau, N. & Erlandsen, H. The metabolic and molecular bases of tetrahydrobiopterin-responsive phenylalanine hydroxylase deficiency. Mol. Genet. Metab. 82, 101–111 (2004).

40. Ge, W. et al. In vivo evaluation of guide-free Cas9-induced safety risks in a pig model. Signal Transduct. Target. Ther. 9, 1–14 (2024).

41. Cunningham, S. C. et al. AAV2/8-mediated correction of OTC deficiency is robust in adult but not neonatal Spf(ash) mice. Mol. Ther. J. Am. Soc. Gene Ther. 17, 1340–1346 (2009).

42. Wang, L. et al. AAV8-mediated Hepatic Gene Transfer in Infant Rhesus Monkeys (Macaca mulatta). Mol. Ther. 19, 2012–2020 (2011).

43. Wang, L., Wang, H., Bell, P., McMenamin, D. & Wilson, J. M. Hepatic Gene Transfer in Neonatal Mice by Adeno-Associated Virus Serotype 8 Vector. Hum. Gene Ther. 23, 533–539 (2012).

44. Suchy, F. P. et al. Genome engineering with Cas9 and AAV repair templates generates frequent concatemeric insertions of viral vectors. Nat. Biotechnol. 1–10 (2024).

45. Turki, A. et al. Minimally invasive (13)C-breath test to examine phenylalanine metabolism in children with phenylketonuria. Mol. Genet. Metab. 115, 78–83 (2015).

46. McCaman, M. W. & Robins, E. Fluorimetric method for the determination of phenylalanine in serum. J. Lab. Clin. Med. 59, 885–890 (1962).

47. Harding, C. O., Wild, K., Chang, D., Messing, A. & Wolff, J. A. Metabolic engineering as therapy for inborn errors of metabolism – development of mice with phenylalanine hydroxylase expression in muscle. Gene Ther. 5, 677–683 (1998).

48. Hoffmann, G., Aramaki, S., Blum-Hoffmann, E., Nyhan, W. L. & Sweetman, L. Quantitative analysis for organic acids in biological samples: batch isolation followed by gas chromatographic-mass spectrometric analysis. Clin. Chem. 35, 587–595 (1989).

