## Supplemental Material for "Enhancement of Therapeutic Transgene Insertion for Murine Phenylketonuria"

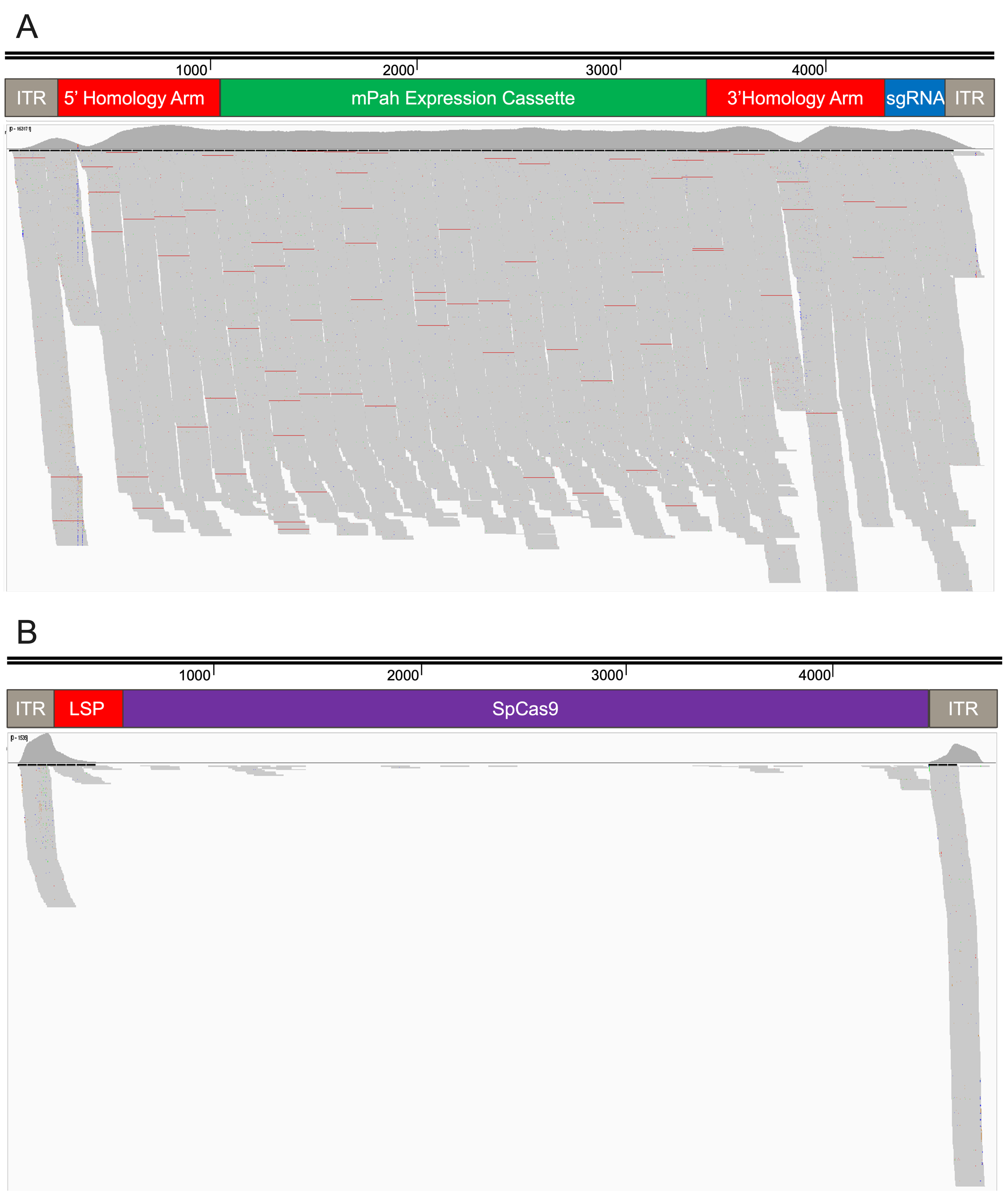

**Figure S1. Genome Annotation Depicting Coverage of Sequencing Reads.** (**A**) Read coverage from enriched genomic DNA sequenced on Illumina next-generation sequencing (NGS) platform and aligned to annotated *Pah* Exon1 repair template. (**B**) Coverage of AAV8 spCas9 vector integrated at genomic target. Histogram indicates read coverage. Gray bars indicate number of aligned reads. Red lines indicate insertions-deletions (indels).

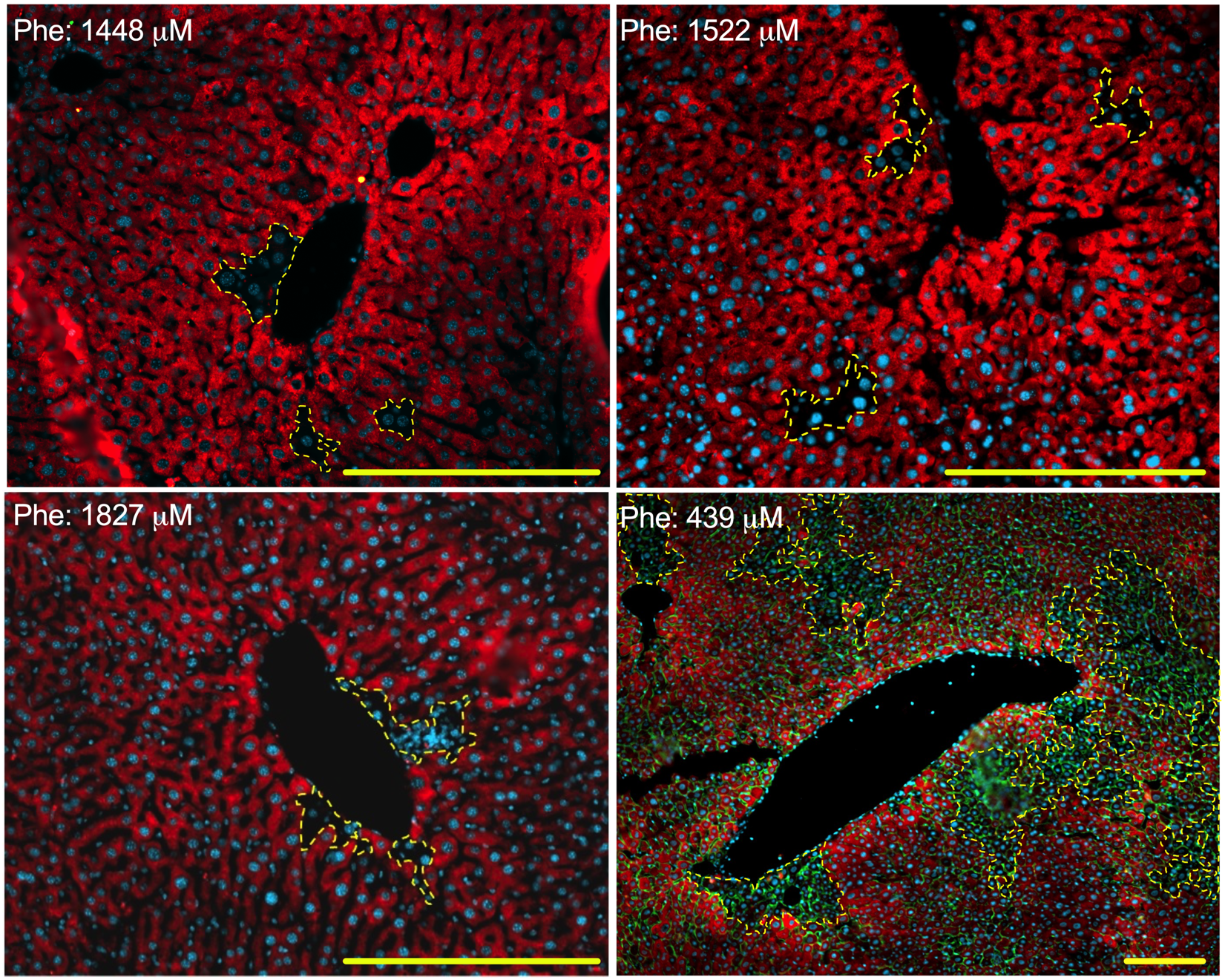

**Figure S2. Immunohistology of Exon7-shCYPOR Hepatocytes.** Anti-CYPOR (Red, cytoplasmic) and anti-lectin (Green, cell membrane) immunostaining of female mice after injection of *Pah* Exon7-shCYPOR repair template and Cas9 AAV8 treatment and 18 weeks on 1.9% APAP diet. Dotted lines surround CYPOR negative colonies. Top, bottom right: 200x magnification; bottom right: 100x magnification. Scale bars, 100 μm.

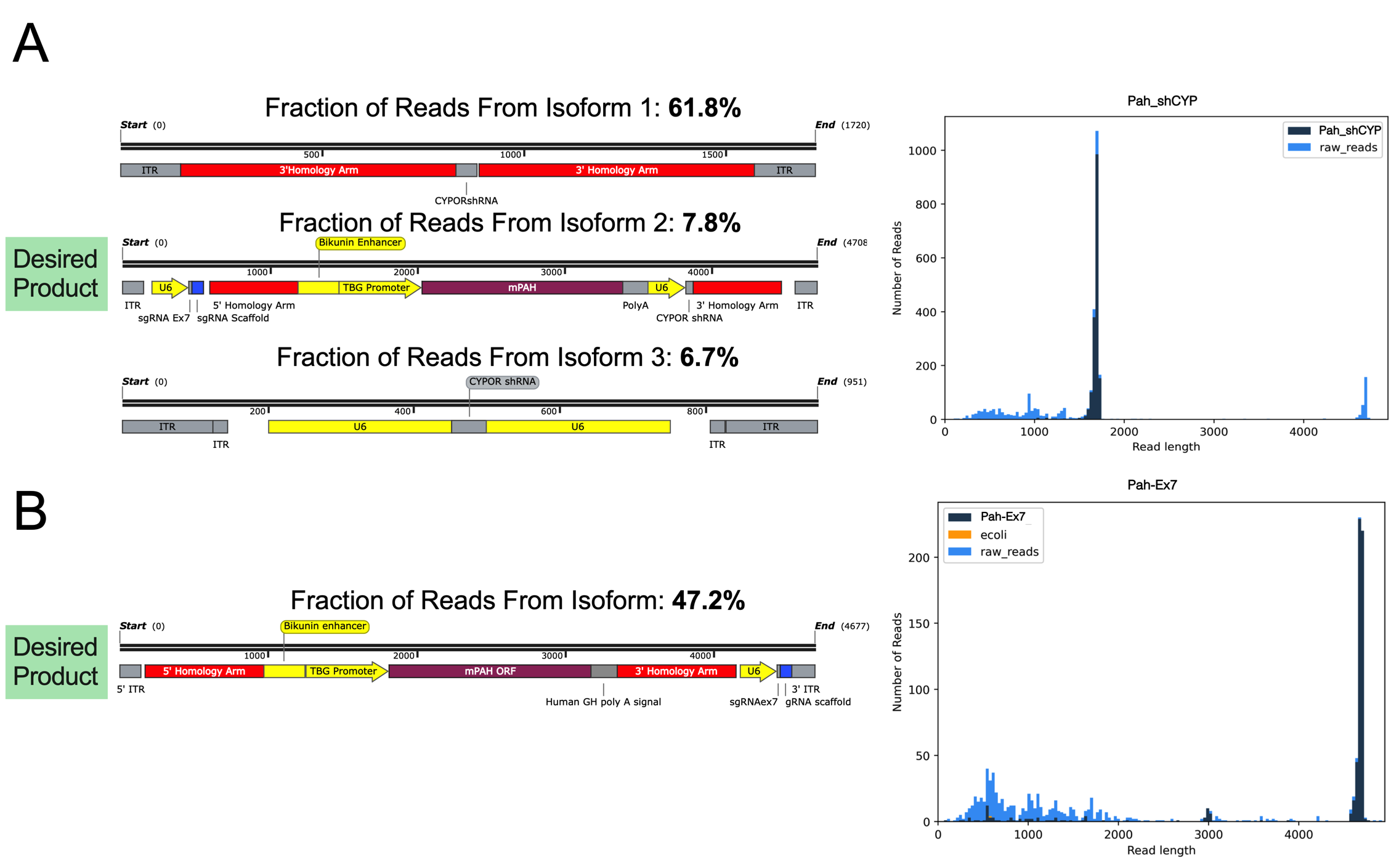

**Figure S3. *Pah* Exon7-shCYPOR Vector Genome Sequencing.** (**A**) Oxford Nanopore sequencing of *Pah* Exon7-shCYPOR AAV8 vector genome. Consensus sequences of three isoforms listed in order of prevalence. Right: Histogram illustrating size of all sequenced reads. (**B**) Oxford Nanopore sequencing of Pah Exon7 AAV8 vector genome with only one consensus isoform sequence. Right: Histogram illustrating size of all sequenced reads.

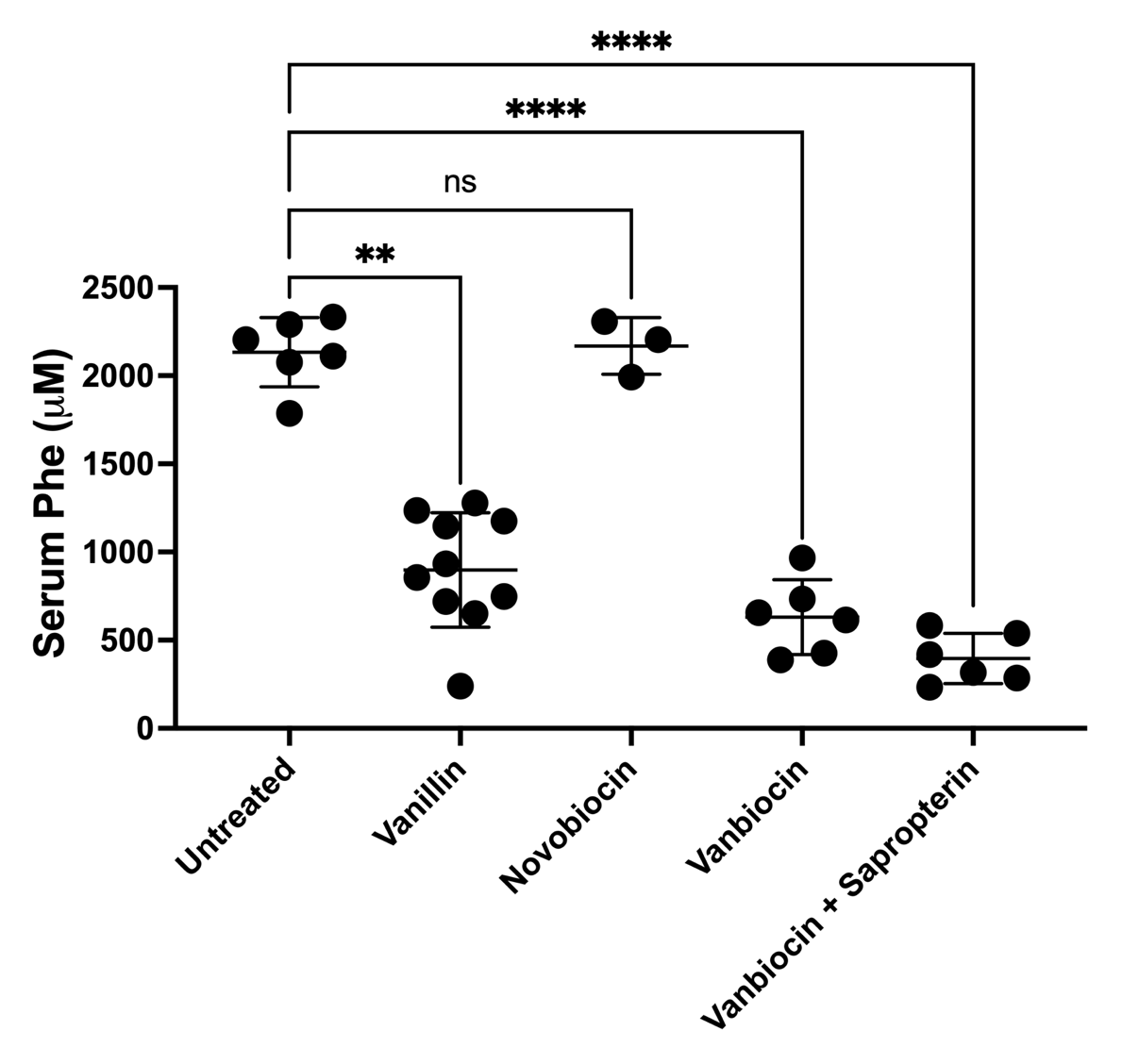

**Figure S4. Combinatorial Effect of Small Molecules on HDR.** Serum phenylalanine concentrations after Cas9/*Pah* Exon7 AAV8 treatment with differing small molecule treatment. *****P* <0.0001, ***P* = 0.0013, and *P* = 0.9869 by one-way ANOVA with Dunnett’s multiple comparison test. Data are means ± SD.

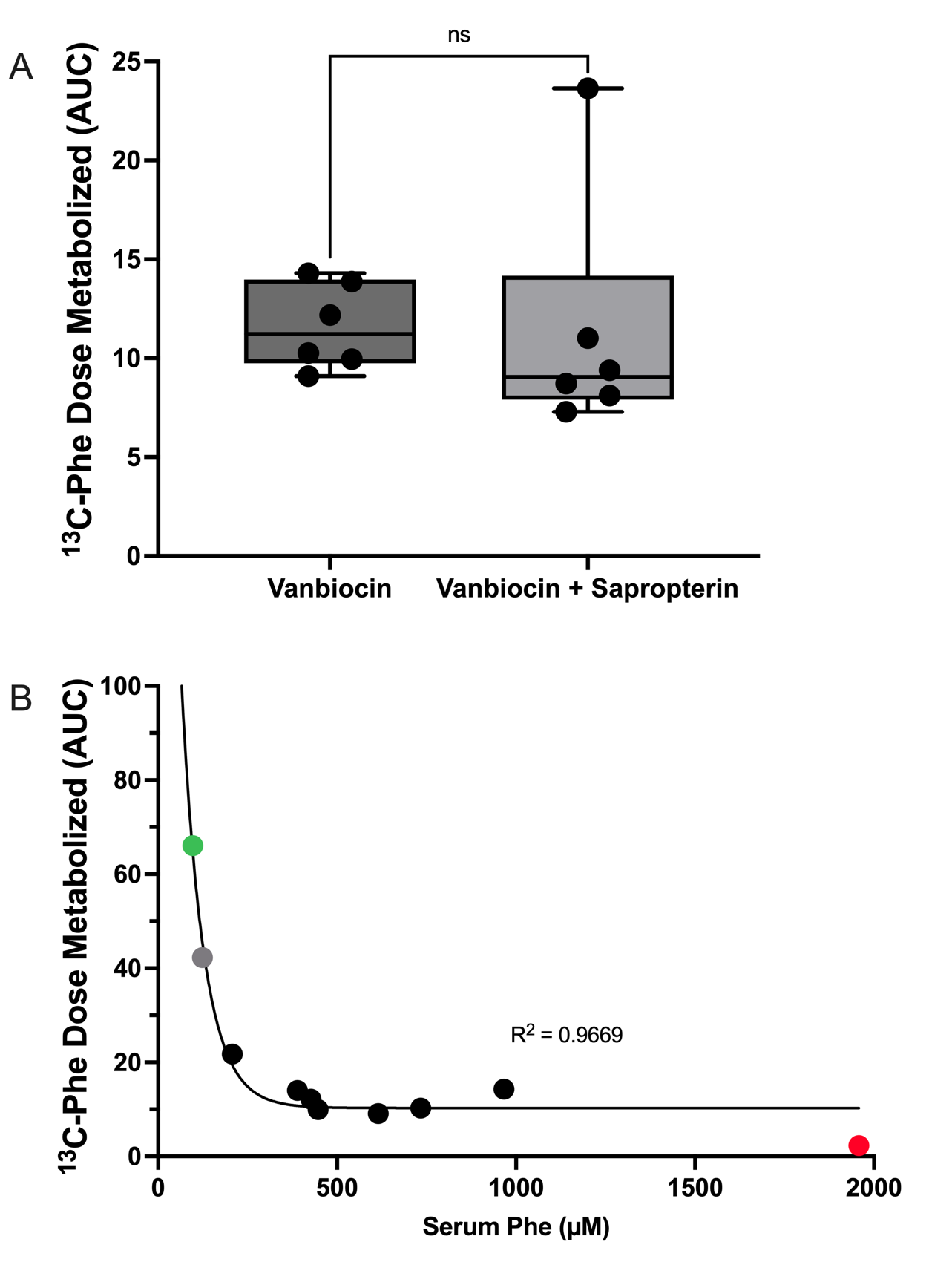

**Figure S5. Stable Isotope Breath Test Data.** (**A**) A stable isotope breath test, measuring the area under the curve (AUC) of exhaled radiolabeled ^13^CO_2_ over 30 minutes following IP administration of 1-^13^C-phenylalanine, was conducted on Vanbiocin-treated mice, with and without sapropterin gavage administered 4 hours prior to the test. *P* = 0.7612 by unpaired T-test. (**B**) Stable isotope breath data of all Vanbiocin treated animals (Y-axis, AUC), including those heterozygous for the *Pah*^Δexon1^ mutation, versus blood phenylalanine concentrations. A single untreated wild-type (Green, Phe = 97 µM), *Pah*^Δexon1/+^ (Grey, Phe = 124 µM), and *Pah*^Δexon1/Δexon1^ (Red, Phe = 1958 µM) mouse were included in this study. Best fit to exponential curve (R^2^ = 0.9655). Data are means ± SD.

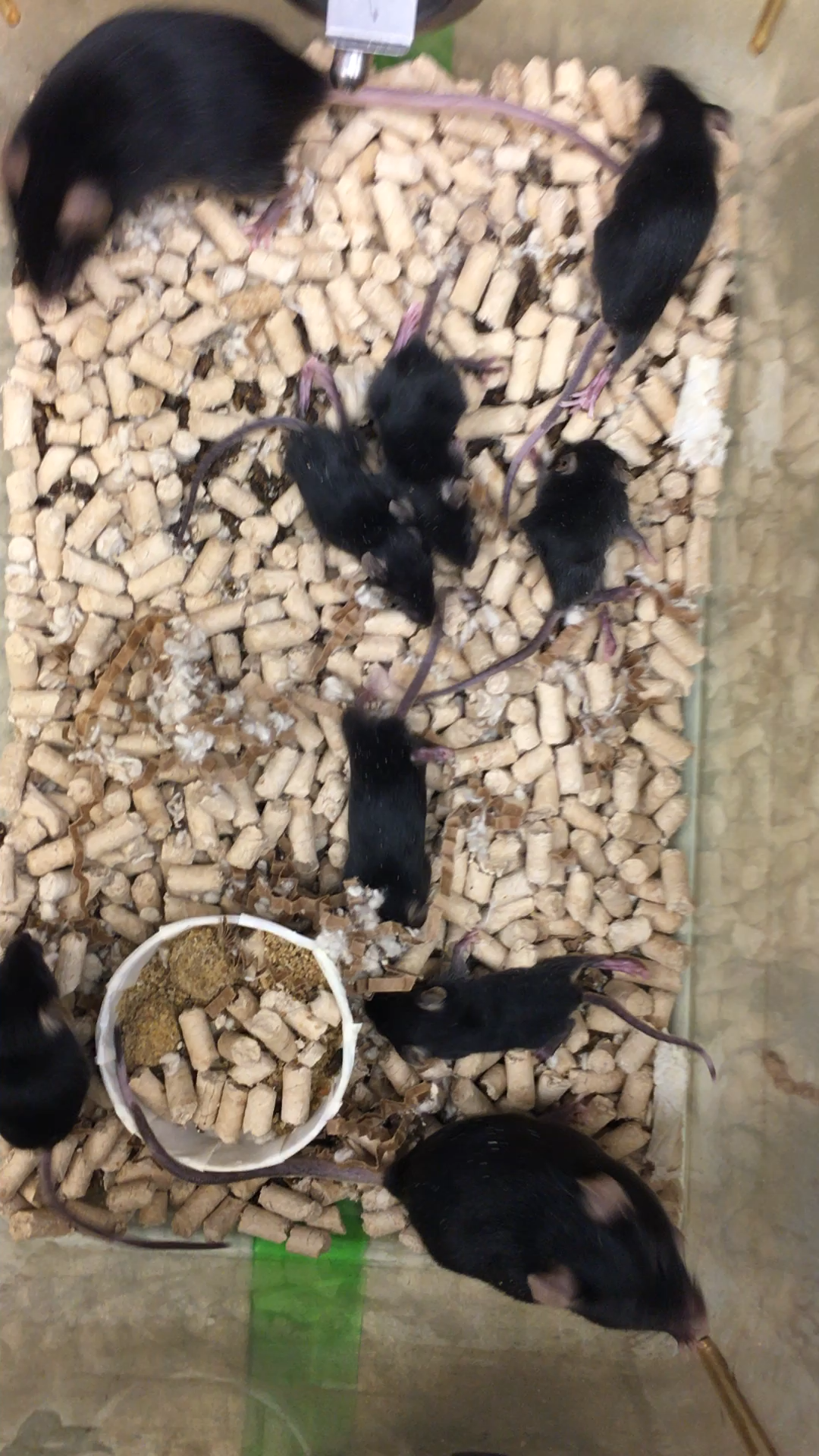

**Supplemental Movie 1.** **Motor dysfunction in AAV8 Pah Dexon7-shCYPOR treated litter.** Mice treated with AAV8 vectors containing *Cypor* shRNA and Cas9, along with IP injections of 100 mg/kg of vanillin, at ~P16.

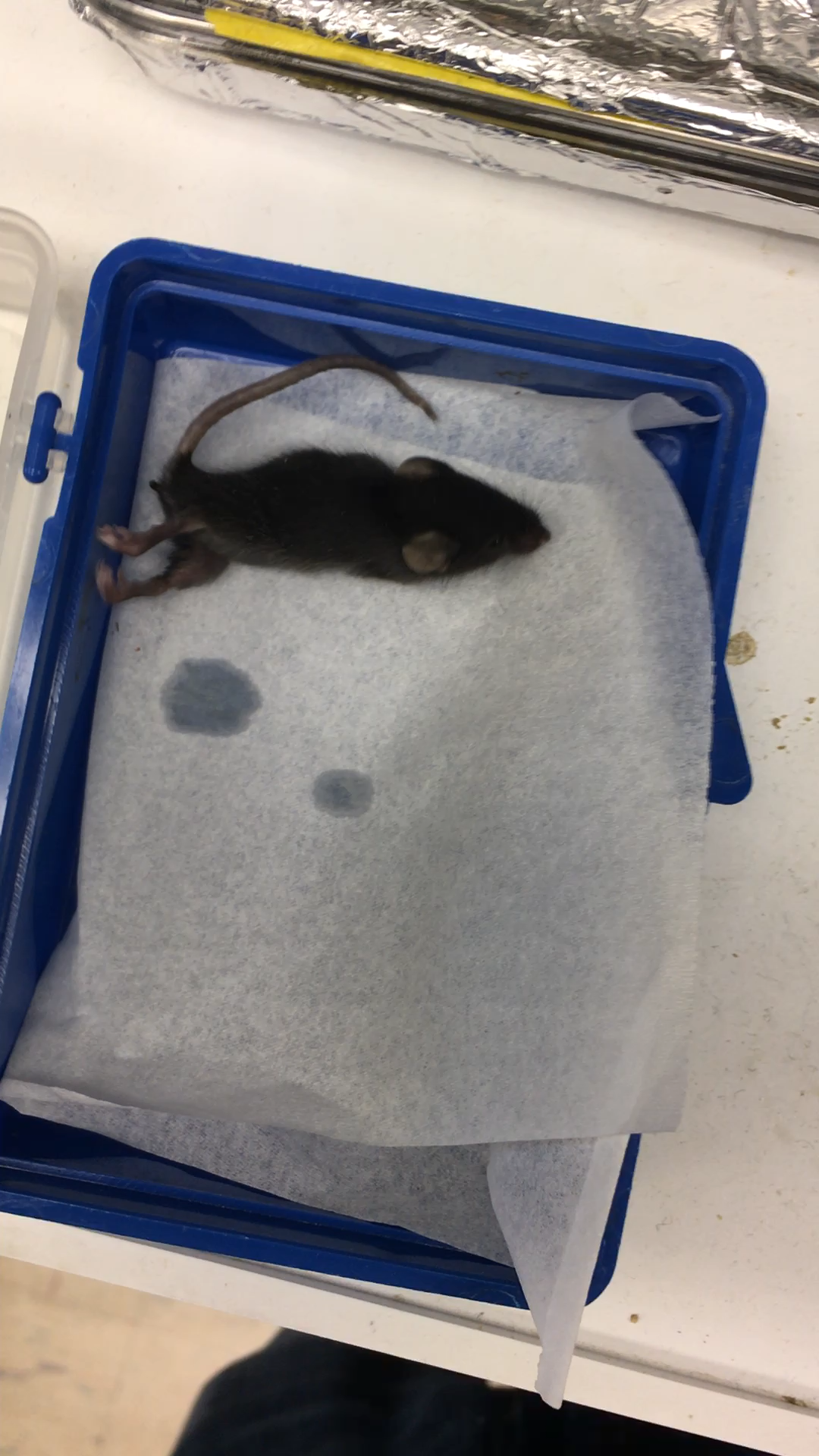

**Supplemental Movie 2. Motor dysfunction in AAV8 Pah Dexon7-shCYPOR treated individual.** Mouse treated with AAV8 vectors containing *Cypor* shRNA and Cas9, along with IP injections of 100 mg/kg of vanillin, at ~P16.
